# Toward a Genome Scale Sequence Specific Dynamic Model of Cell-Free Protein Synthesis in *Escherichia coli*

**DOI:** 10.1101/215012

**Authors:** Nicholas Horvath, Michael Vilkhovoy, Joseph A. Wayman, Kara Calhoun, James Swartz, Jeffrey D. Varner

## Abstract

Cell-free protein expression systems have become widely used in systems and synthetic biology. In this study, we developed an ensemble of dynamic *E. coli* cell-free protein synthesis (CFPS) models. Model parameters were estimated from a training dataset for the cell-free production of a protein product, chloramphenicol acetyltransferase (CAT). The dataset consisted of measurements of glucose, organic acids, energy species, amino acids, and CAT. The ensemble accurately predicted these measurements, especially those of the central carbon metabolism. We then used the trained model to evaluate the optimality of protein production. CAT was produced with an energy efficiency of 12%, suggesting that the process could be further optimized. Reaction group knockouts showed that protein productivity and the metabolism as a whole depend most on oxidative phosphorylation and glycolysis and gluconeogenesis. Amino acid biosynthesis is also important for productivity, while the overflow metabolism and TCA cycle affect the overall system state. In addition, the translation rate is shown to be more important to productivity than the transcription rate. Finally, CAT production was robust to allosteric control, as was most of the network, with the exception of the organic acids in central carbon metabolism. This study is the first to use kinetic modeling to predict dynamic protein production in a cell-free *E. coli* system, and should provide a foundation for genome scale, dynamic modeling of cell-free *E. coli* protein synthesis.

## Introduction

Cell-free protein expression has become a widely used research tool in systems and synthetic biology, and a promising technology for personalized point of use biotechnology [1]. Cell-free systems offer many advantages for the study, manipulation and modeling of metabolism compared to *in vivo* processes. Central amongst these, is direct access to metabolites and the biosynthetic machinery without the interference of a cell wall, or complications associated with cell growth. This allows us to interrogate (and potentially manipulate) the chemical microenvironment while the biosynthetic machinery is operating, potentially at a fine time resolution. Cell-free protein synthesis (CFPS) systems are arguably the most prominent examples of cell-free systems used today [2]. However, CFPS is not new; CFPS in crude *E. coli* extracts has been used since the 1960s to explore fundamental biological mechanisms. For example, Matthaei and Nirenberg used *E. coli* cell-free extract in ground-breaking experiments to decipher the sequencing of the genetic code [3, 4]. Spirin and coworkers later improved protein production in cell free extracts by continuously exchanging reactants and products; however, while these extracts could run for tens of hours, they could only synthesize a single product and were energy limited [5]. More recently, energy and cofactor regeneration in CFPS has been significantly improved; for example ATP can be regenerated using substrate level phosphorylation [6] or even oxidative phosphorylation [2]. Today, cell-free systems are used in a variety of applications ranging from therapeutic protein production [7] to synthetic biology [8, 1]. Moreover, there are also several CFPS technology platforms, such as the PANOx-SP and Cytomin platforms developed by Swartz and coworkers [9, 2], and the TX/TL platform of Noireaux [10]. However, if CFPS is to become a mainstream technology for applications such as point of care biomanufacturing, we must first understand the performance limits of these systems, and eventually optimize their yield and productivity. A critical tool towards this goal is the development of a CFPS mathematical model.

Mathematical modeling has long contributed to our understanding of metabolism [11]. Decades before the genomics revolution, mechanistically structured metabolic models arose from the desire to predict microbial phe-notypes resulting from changes in intracellular or extracellular states [12]. The single cell *E. coli* models of Shuler and coworkers pioneered the con-struction of large-scale, dynamic metabolic models that incorporated multiple regulated catabolic and anabolic pathways constrained by experimentally determined kinetic parameters [13]. Shuler and coworkers generated many single cell kinetic models, including single cell models of eukaryotes [14, 15], minimal cell architectures [16], and DNA sequence based whole-cell models of *E. coli* [17]. More recent studies have extended the approach to integrate disparate models of cellular processes in *M. genitalium* [18], describe dozens of mutant strains in *E. coli* with a single kinetic model [19], and identify industrially useful target enzymes in *E. coli* to improve 1,4-butanediol production [20]. However, cell-free genome scale kinetic models of industrially important organisms such as *E. coli* have yet to be constructed.

In this study, we developed an ensemble of kinetic cell-free protein synthesis (CFPS) models using dynamic metabolite measurements from an early glucose powered Cytomin *E. coli* cell-free extract. While cell-free technology has evolved considerably since these measurements were taken, developing a model using a previous generation CFPS platform offers several unique advantages. First and foremost, is the ability to directly compare the dif-ferent improvements established by purely experimental means, to those estimated from a mathematical model. The CFPS model equations were formulated using the hybrid cell-free modeling framework of Wayman et al. [21], which integrates traditional kinetic modeling with a logical rule-based description of allosteric regulation. Model parameters were estimated from measurements of glucose, organic acids, energy species, amino acids, and the protein product, chloramphenicol acetyltransferase (CAT) over the course of a three hour protein synthesis reaction. A constrained Markov Chain Monte Carlo (MCMC) approach was used to minimize the squared difference between model simulations and experimental measurements, where a plausible range for each kinetic parameter was established from BioNumbers [22]. The ensemble of parameter sets described the training data with a median cost greater than two orders of magnitude smaller than a population of random parameter sets constructed using the same literature parameter constraints. We then used the ensemble of kinetic models to analyze the performance of the CFPS system, and to estimate the pathways most important to protein production. We calculated that CAT was produced with an energy efficiency of 12%, suggesting that much of the energy resources for protein synthesis were diverted to non-productive pathways. By knocking out metabolic enzymes in groups, we showed that metabolism and protein production in particular depended upon oxidative phosphorylation and glycolysis /gluconeogenesis. Taken together, this study provides a foundation for sequence specific genome scale, dynamic modeling of cell-free *E. coli* protein synthesis.

## Results

The cell-free *E. coli* metabolic network was constructed by removing growth associated reactions from the *i*AF1260 reconstruction of K-12 MG1655 *E. coli* [23], and by adding reactions describing chloramphenicol acetyltransferase (CAT) biosynthesis (Fig. 1). In addition, reactions that were knocked out in the host strain used to prepare the extract were removed from the network (ΔspeA, ΔtnaA, ΔsdaA, ΔsdaB, ΔgshA, ΔtonA, ΔendA). Lastly, we added the transcription and translation template reactions of Allen and Palsson for the specific proteins of interest [24]. The metabolic network, which contained XX metabolites and YY reactions, is available in the supplemental materials. The dynamic CFPS model equations were formulated using the hybrid cell-free modeling framework of Wayman et al. [21]. An ensemble of model parameter sets (N = 3,000) was estimated from measurements of glucose, CAT, organic acids (pyruvate, lactate, acetate, succinate, malate), energy species (A(x)P, G(x)P, C(x)P, U(x)P), and 18 of the 20 proteinogenic amino acids [25] using a constrained Markov Chain Monte Carlo (MCMC) approach.

**Figure 1:**
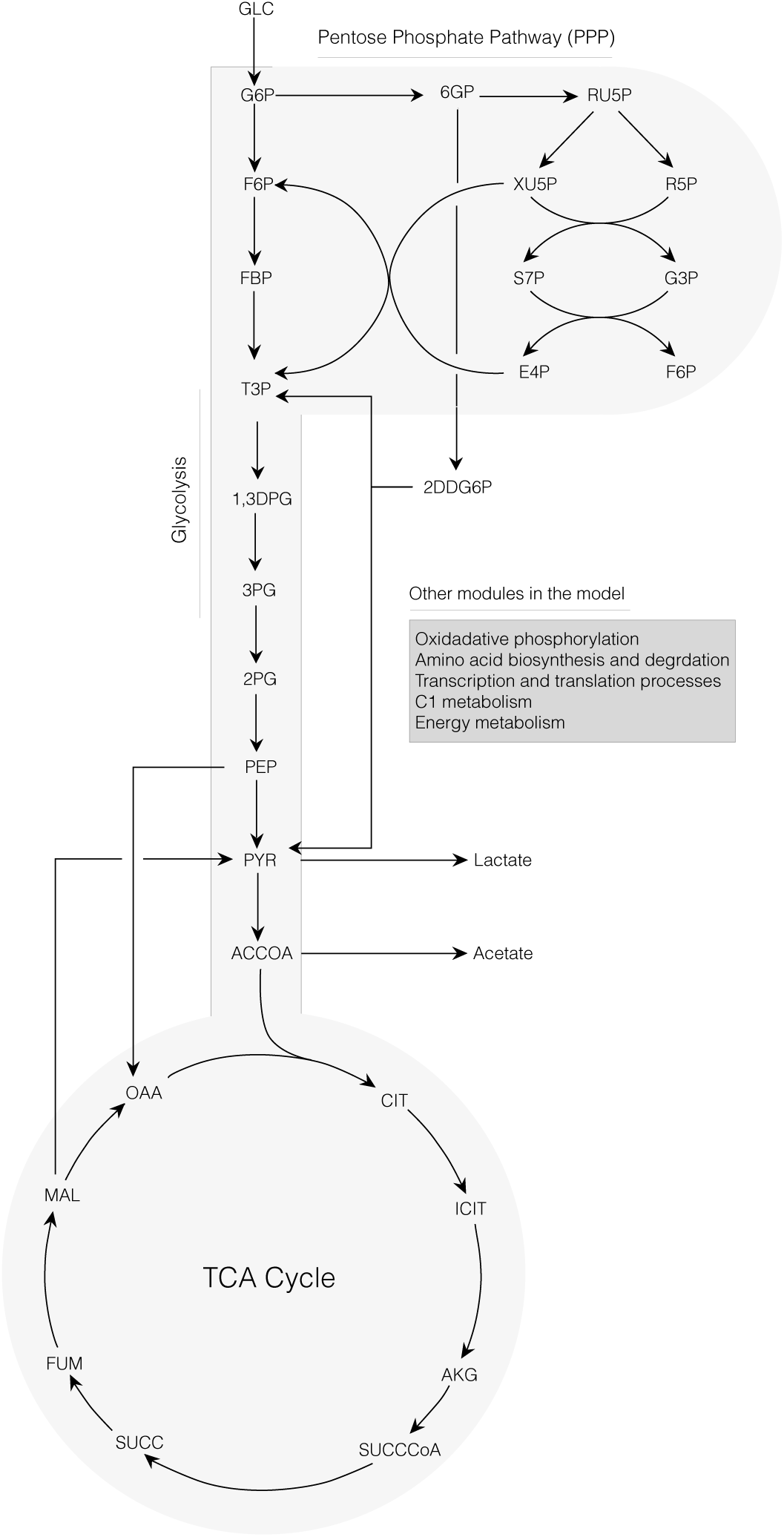
Schematic of the core portion of the cell-free *E. coli* metabolic network. Metabolites of glycolysis, pentose phosphate pathway, Entner-Doudoroff pathway, and TCA cycle are shown. Metabolites of oxidative phosphorylation, amino acid biosynthesis and degradation, transcription/translation, chorismate metabolism, and energy metabolism are not shown.

The MCMC algorithm minimized the squared difference (residual) between the training data and model simulations starting from an initial parameter set assembled from literature and inspection. Bounds on permissible parameter values were established using studies from the BioNumbers database [22]. For each newly generated parameter set, we re-solved the bal-ance equations and calculated the cost function; all sets with a lower cost (and some with higher cost) were accepted into the ensemble. Parameter sets were also required to meet strict ordinary differential equation solver tolerances, to ensure numerical stability. Approximately N = 3,000 sets were accepted into an initial ensemble; N = 100 sets were then selected based upon error for the final ensemble. The final ensemble had a mean Pearson correlation coefficient of 0.78; this suggested parameter sets were not over sampled in the region of a local minimum. The median maximum reaction rate (*V*_*max*_) across the ensemble was 11.6 mM/h, assuming a total cell-free enzyme concentration of 170 nM. This *V_max_* corresponded to a median catalytic rate of 19 s^−1^ across the ensemble; this was in agreement with the 13.7 s^−1^ median catalytic rate found by Bar-Even and coworkers [26]. The median enzyme activity decay constant was 0.0045 h^−1^, corresponding to an enzyme activity half life of 6 days. The median saturation constant was 1.0 mM; this is within one order of magnitude of the 130 *μ*M reported by Bar-Even and coworkers. Lastly, both the median control gain parameter, and the control order parameter in the ensemble were order 1. While the maximum reaction rates of the ensemble were distributed evenly across the allowed range (Fig. S1A), the saturation constants were clustered around the upper and lower bounds (Fig. S1B).

The ensemble of kinetic CFPS models captured the time evolution of protein biosynthesis, and the consumption and production of organic acid, amino acid and energy species. Central carbon metabolites (Fig. 2, top), energy species (Fig. 4), and amino acids (Fig. 3) were captured by the ensemble and the best-fit set. The constrained MCMC approach estimated parameter sets with a median error more than two orders of magnitude less than random parameter sets generated within the same parameter bounds established from literature (Fig. 5); thus, we have confidence in the predictive capability of the estimated parameters. For 29 of the 37 measurements in the training dataset, the mean Akaike information criterion (AIC) of the ensemble was lower than that of the random sets, signifying a better fit of the data (Table 3). For the other 8 measurements, the random AIC was lower than the ensemble by an amount less than the standard deviation of either the random AIC or ensemble AIC (with the exception of isoleucine, which was quite close: 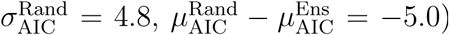. Taken together, these results suggested that the parameter ensemble modeled cell free metabolism and protein production, significantly better than if sampled randomly, not just overall but for the majority of individual measurements.

**Figure 2:**
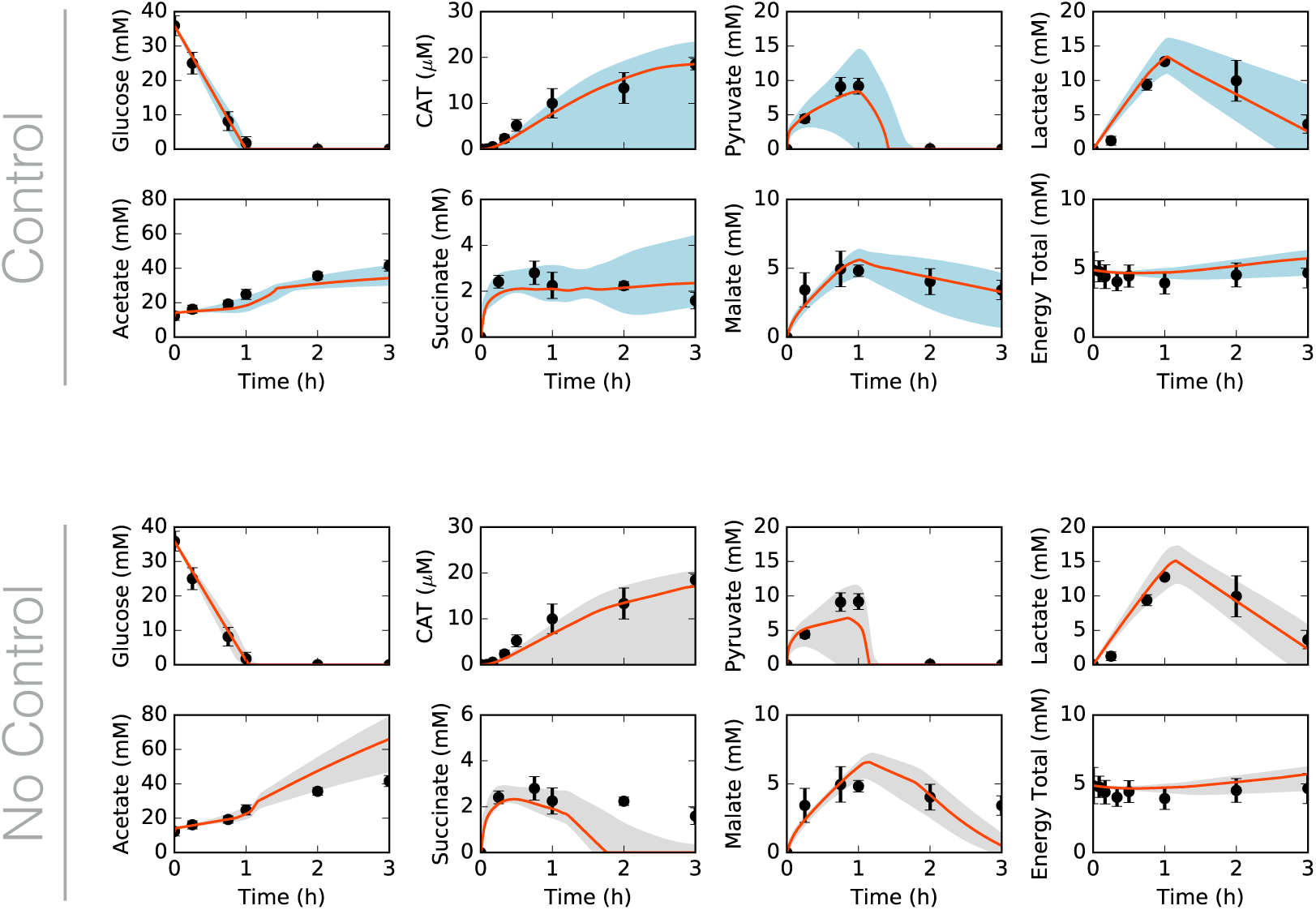
Central carbon metabolism in the presence (top) and absence (bottom) of allosteric control, including glucose (substrate), CAT (product), and intermediates, as well as total concentration of energy species. Best-fit parameter set (orange line) versus experimental data (points). 95% confidence interval (blue or gray shaded region) over the ensemble of 100 sets.

**Figure 3:**
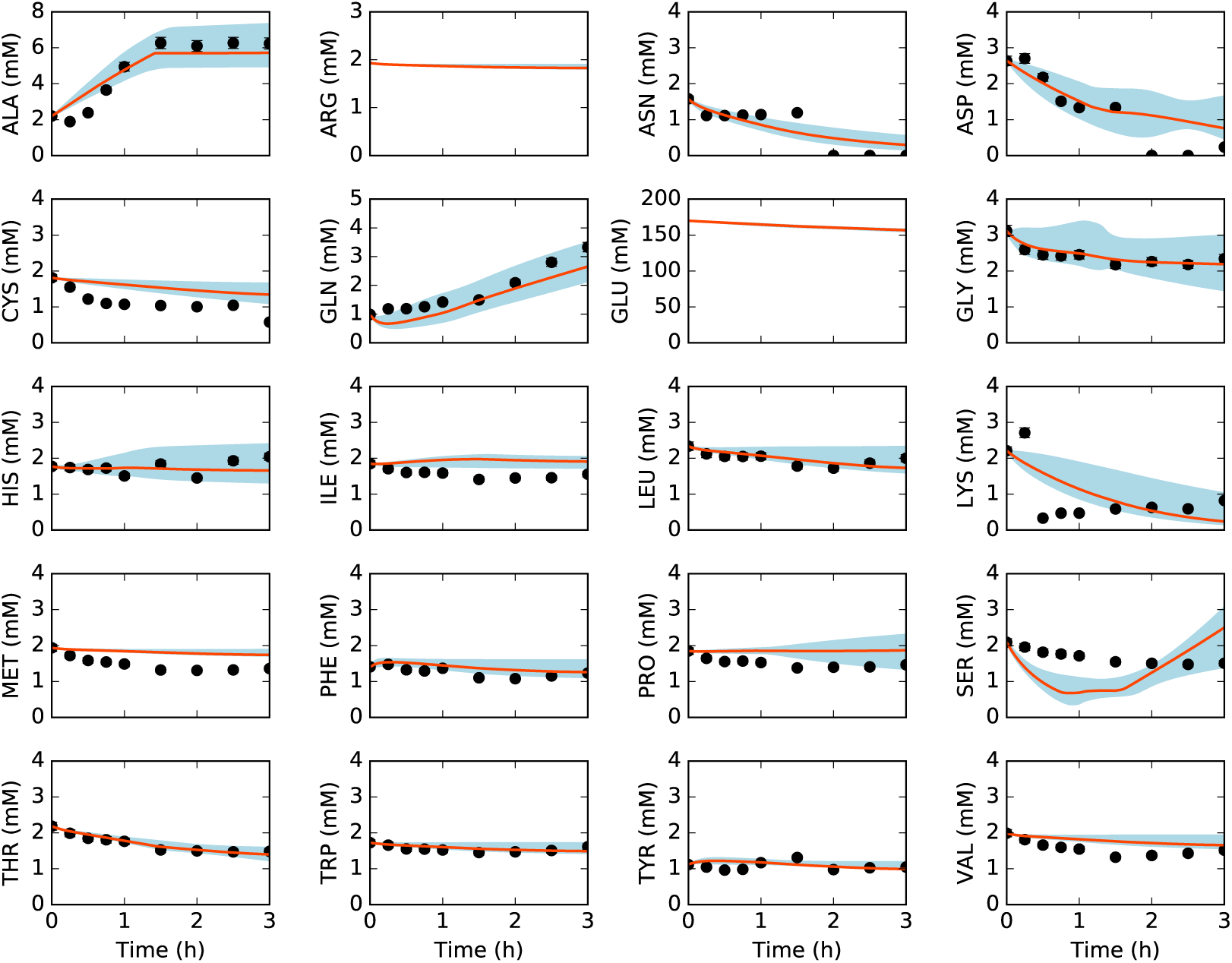
Amino acids in the presence of allosteric control. Best-fit parameter set (orange line) versus experimental data (points). 95% confidence interval (blue shaded region) over the ensemble of 100 sets.

**Figure 4:**
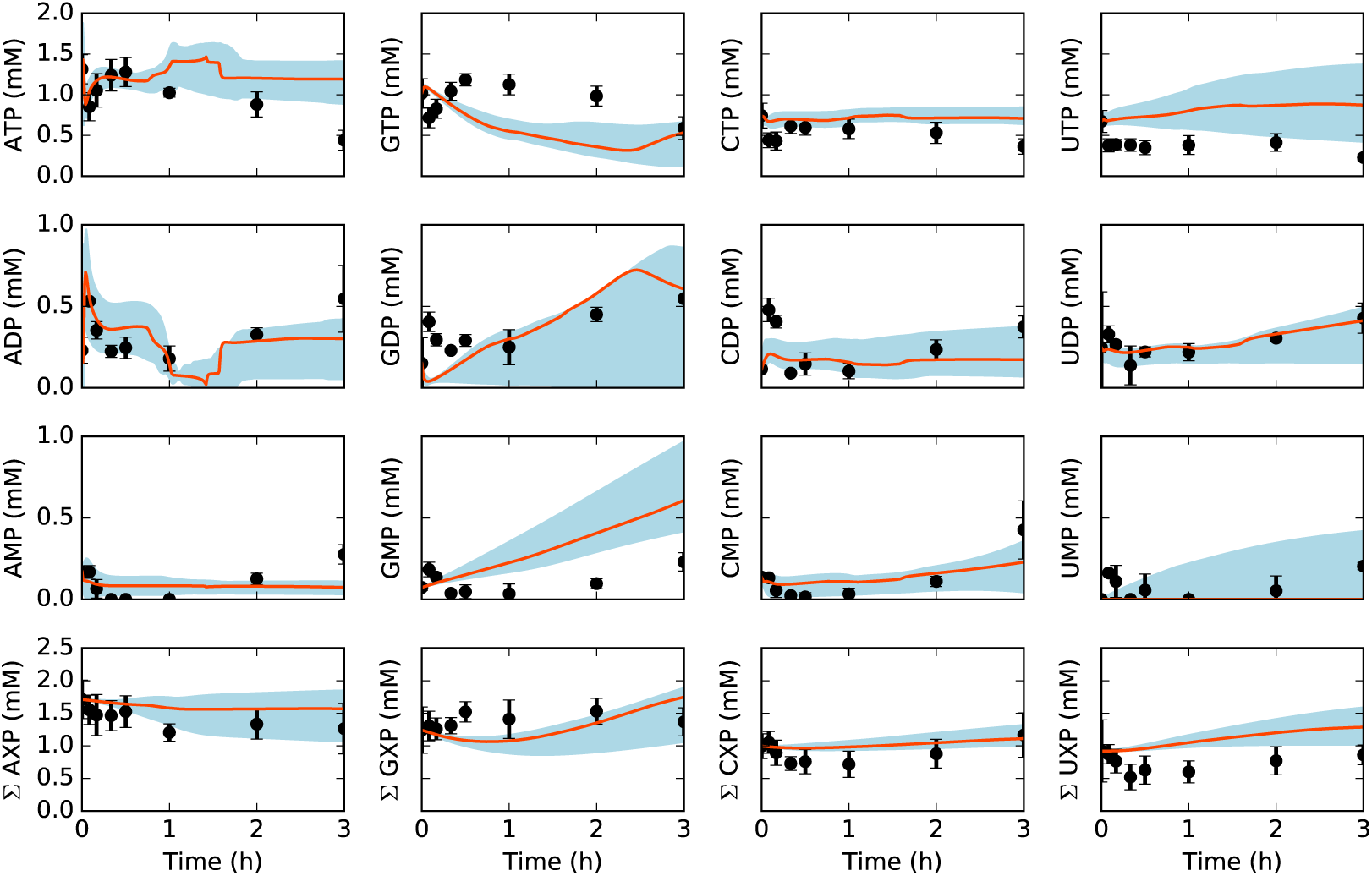
Energy species and energy totals by base in the presence of allosteric control. Best-fit parameter set (orange line) versus experimental data (points). 95% confidence interval (blue shaded region) over the ensemble of 100 sets.

**Figure 5:**
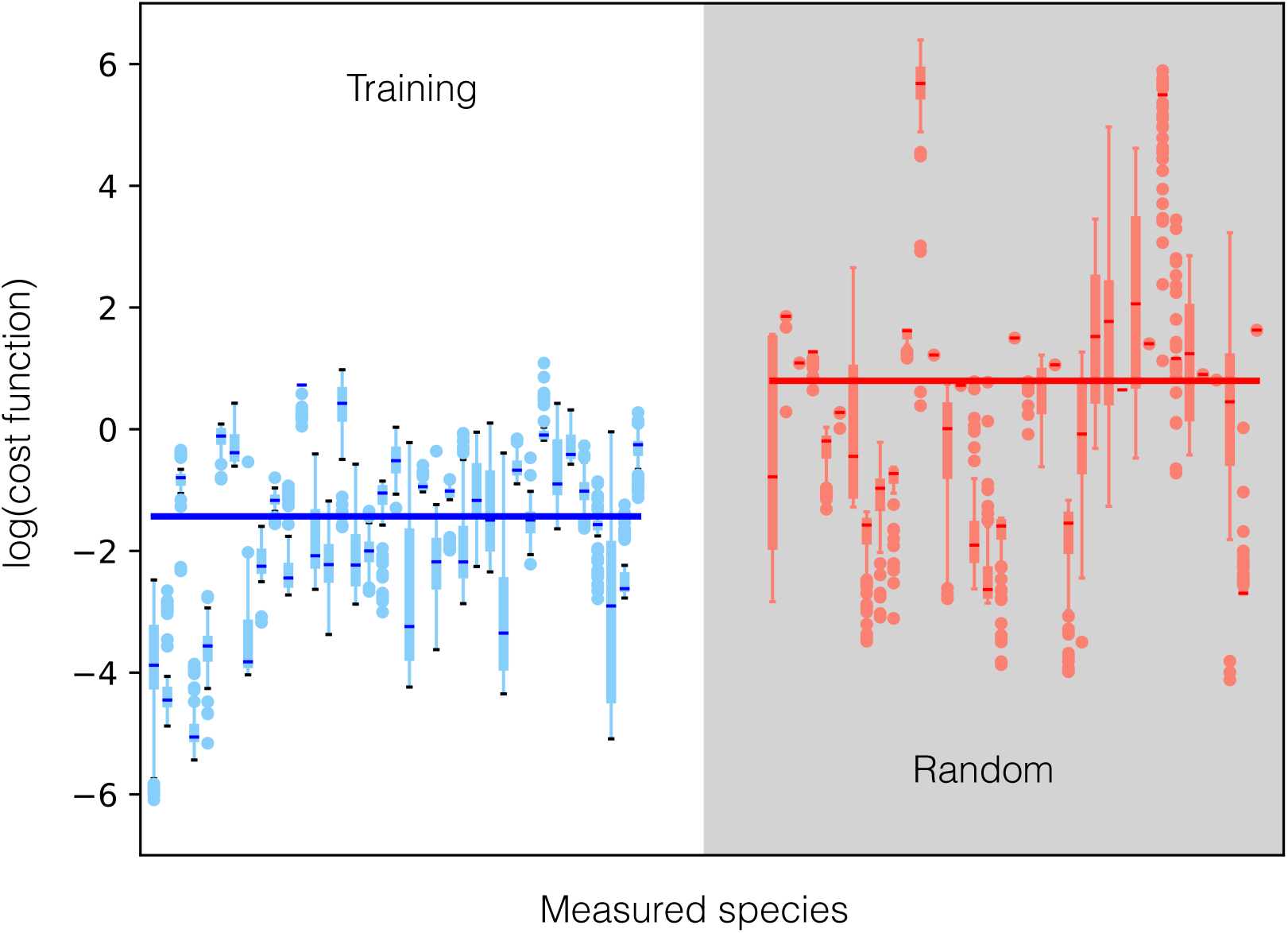
Log of cost function (residual between training data and model simulations) across 37 datasets for data-trained ensemble (blue) and randomly generated ensemble (red, gray background). Median (bars), interquartile range (boxes), range excluding outliers (thin lines), and outliers (circles) for each dataset. Median across all datasets (large bar overlaid).

The model captured the biphasic time course of CAT production. During the first hour glucose powered protein production, and CAT was produced at 8 *μ*M/h; subsequently, pyruvate and lactate reserves were consumed to power metabolism, and CAT was produced less quickly at 5 *μ*M/h. Allosteric control was important to central carbon metabolism, especially pyruvate, acetate, and succinate (Fig. 2, bottom). The difference between the allosteric control and no-control cases was mostly seen in the second phase of CAT production, following glucose exhaustion. Specifically, pyruvate, succinate, and malate consumption and acetate accumulation increased following glucose exhaustion without the allosteric control mechanisms. The rate of acetate accumulation increased by 172%, while the rates of malate, pyruvate, and lactate consumption increased by 146%, 82%, and 9%, respectively. Succinate went from accumulating slightly in the second phase, in the presence of allosteric control, to being fully consumed. However, CAT production was robust to the removal of allosteric control, as seen in both the fits against data and the metabolic fluxes (see supplementary information). While ATP generation varied when allosteric control was removed, ATP expenditure toward CAT production did not. Most of the fluxes that differed between the two cases involved PEP and pyruvate, which directly participated in many of the reactions modulated by allosteric control. Taken together, the ensemble of kinetic models was consistent with time series measurements of the cell free production of a model protein. Although the ensemble described the experimental data, it was unclear which kinetic parameters and pathways most influenced CAT production. To explore this question, we performed reaction group knockout analysis.

The importance of CFPS pathways was estimated using pathway group knockout analysis (Fig. 7). The metabolic network was divided into 19 reaction groups, spanning central carbon metabolism, energetics, and amino acid biosynthesis. The response in the productivity or overall system state was calculated for single or pairwise deletion of each of these reaction groups. Lastly, the overall effect of the deletion of a pathway was estimated by sum-ming the single and pairwise effects (summation across the columns of the response array). Glycolysis/gluconeogenesis and oxidative phosphorylation had the greatest effect on both productivity and system state. This supports previous studies that have suggested oxidative phosphorylation is occurring in a cell-free system [2]; Jewett and coworkers observed a decrease in CAT yield, ranging from 1.5-fold to 4-fold, when inhibiting oxidative phosphorylation reactions in the Cytomim cell-free platform, using both pyruvate and glutamate as substrates. CAT productivity was also affected by two sectors of amino acid biosynthesis: alanine/aspartate/asparagine, and glutamate/ glutamine biosynthesis. This was consistent with aspartate, glutamate, and glutamine being key reactants in the biosynthesis of many other amino acids, all of which are required for CAT synthesis. Meanwhile, the TCA cycle and overflow metabolism (which included acetyl-coA/acetate reactions and the interconversion of pyruvate and lactate) also had a significant effect on the system state. These reactions directly impacted key system species: succinate and malate in the TCA cycle, and acetate, pyruvate, and lactate in the overflow metabolism. In addition, the relative influence of transcription and translation were interrogated via Sobol sampling [27]. Productivity was seen to have a sensitivity of 0.43 ± 0.06 with respect to the maximum reaction rate of transcription, and 0.66 ± 0.08 for the maximum reaction rate of translation. Thus, translation was the limiting step of cell free protein synthesis.

**Figure 6:**
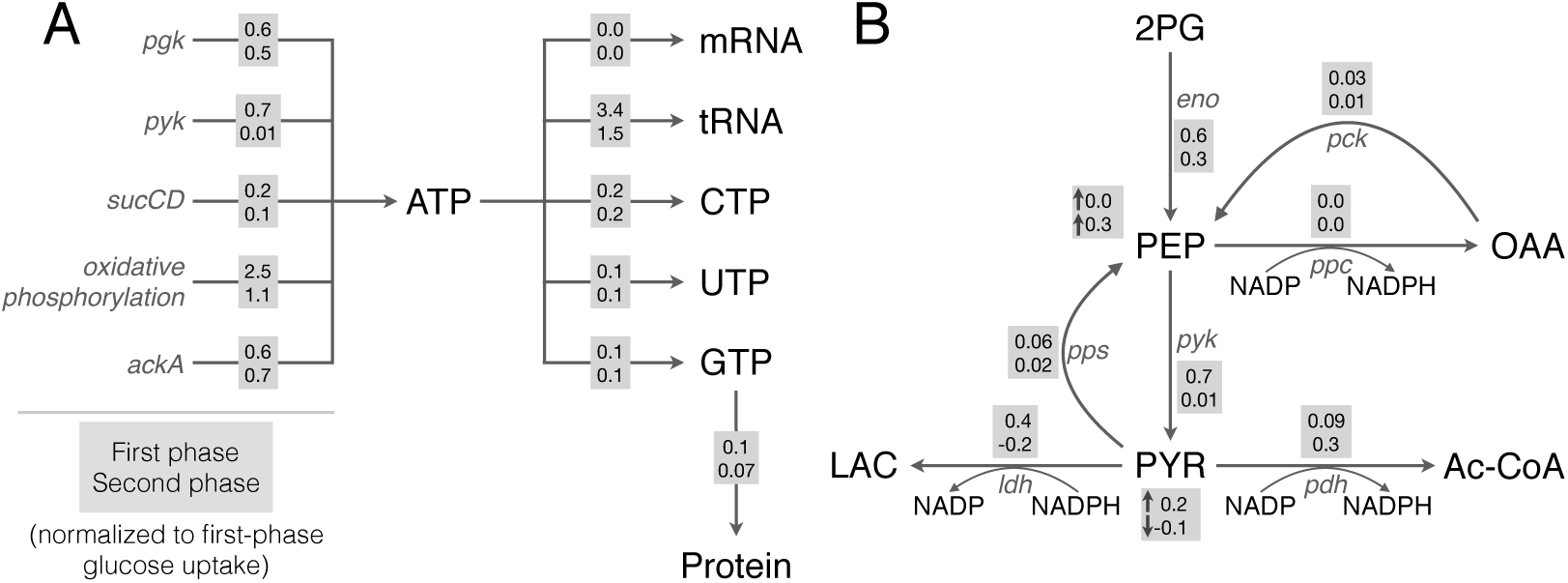
Key reaction fluxes of the network, in the first (gray boxes, top row) and second (gray boxes, bottom row) phases of metabolism. A. Fluxes of ATP generation and consumption, and GTP consumption toward protein synthesis. B. Fluxes of glycolysis and lactate and acetate metabolism. Fluxes are normalized to the first-phase glucose uptake rate. For PEP and pyruvate, accumulation (normalized to glucose uptake) is also shown.

**Figure 7:**
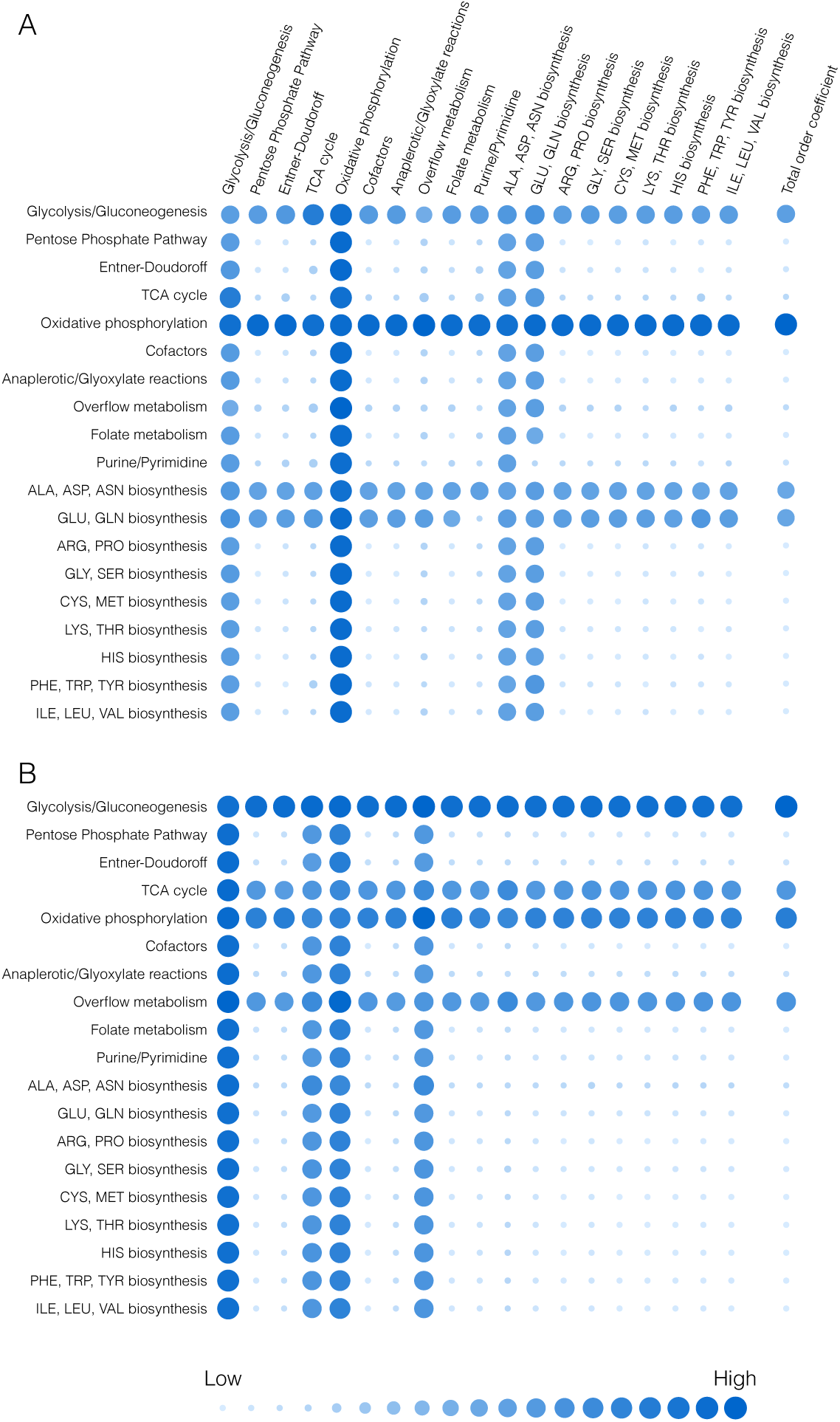
Effect of group knockouts on system. A. Change in CAT productivity when one (diagonal) or two (off-diagonal) reaction groups are turned off. B. Change in system state (only species for which data exist) when one (diagonal) or two (off-diagonal) reaction groups are turned off. Total-order effect for each group calculated as the sum of first-order effect and all pairwise effects. Larger and darker circles represent greater effects.

The energy efficiency of CAT production, as well as the sources of energy generation and consumption, were tracked for the best-fit set. Energy efficiency was calculated as the ratio of transcription and translation rates (weighted by the associated ATP costs of each step) to the amount of ATP generated by all sources. During the first phase of protein production, with glucose as the substrate, CAT was produced with a productivity of 8 *μ*M/h and an energy efficiency of 10%. Oxidative phosphorylation accounted for greater than 50% of the ATP generated during the rapid phase of protein production (Table 1). The organic acids that accumulated in the first phase (with the exception of acetate) were then utilized as substrates in the second phase, once glucose was depleted. We assumed the second phase of CAT production was powered largely by pyruvate; although malate was consumed in the second phase, it only accounted for 11% of substrate consumption. Furthermore, lactate is connected in the stoichiometry only to pyruvate. Thus, it is reasonable to consider the second phase as pyruvate-driven production. Interestingly, while this mode of protein production was slower (5 *μ*M/h), it exhibited a higher energy efficiency (14%). Of the ATP generated, about half was observed to come from oxidative phosphorylation in each of the two phases of production (Table 1, R_atp). Another 30% was generated by glycolysis during the first phase (R_pgk,R_pyk), which decreased to approximately 20% following glucose exhaustion. However, glycolysis was also amongst the largest consumers of ATP during first phase of production (R_glk_atp, R_pfk) (Table 2). The TCA cycle (R_sucCD) contributed 3% of to the overall ATP generation in the first phase and 5% in the second. The hypothesis that pyruvate drives the second phase explains this; stores of accumulated pyruvate can be converted to acetyl-CoA, as well as OAA (via PEP), and thus power the TCA cycle just as when glucose was available. Interestingly, ATP generation through acetate metabolism (R_ackA) increased from 12% in the first phase to 28% in the second. Amino acid degradation also contributes a negligible amount to energy production. While the efficiency of production was higher for the pyruvate-driven phase, it was still relatively low, suggesting that there is room for platform optimization. Taken together, this strengthens the importance of glycolysis and oxidative phosphorylation, and presents a trade-off between productivity and energy efficiency in CFPS.

**Table 1:** Breakdown of ATP generation. Flux through ATP-generating pathways in the first and second phases as percentages of total ATP generation in that phase.

| **Name** | **Index** | **Reaction** | **Phase 1** | **Phase 2** |
| --- | --- | --- | --- | --- |
| R_pgk | 12 | $13\text{DPG} + \text{ADP} \rightarrow 3\text{PG} + \text{ATP}$ | 14% | 21% |
| R_pyk | 18 | $\text{ADP} + \text{PEP} \rightarrow \text{ATP} + \text{PYR}$ | 16% | <1% |
| R_sucCD | 45 | $\text{ADP} + \text{P}_i + \text{SUCCOA} \rightarrow \text{ATP} + \text{COA} + \text{SUCC}$ | 3% | 5% |
| R_atp | 55 | $\text{ADP} + \text{P}_i + 4 \text{H}_e \rightarrow \text{ATP} + 4 \text{H} + \text{H}_2\text{O}$ | 54% | 46% |
| R_ackA | 68 | $\text{ACTP} + \text{ADP} \rightarrow \text{AC} + \text{ATP}$ | 12% | 28% |
| R_asn_deg | 102 | $\text{ASN} + \text{AMP} + \text{PP}_i \rightarrow \text{NH}_3 + \text{ASP} + \text{ATP}$ | <1% | <1% |
| R_thr_deg3 | 109 | $\text{THR} + \text{P}_i + \text{ADP} \rightarrow \text{NH}_3 + \text{FOR} + \text{ATP} + \text{PROP}$ | <1% | <1% |

**Table 2:**
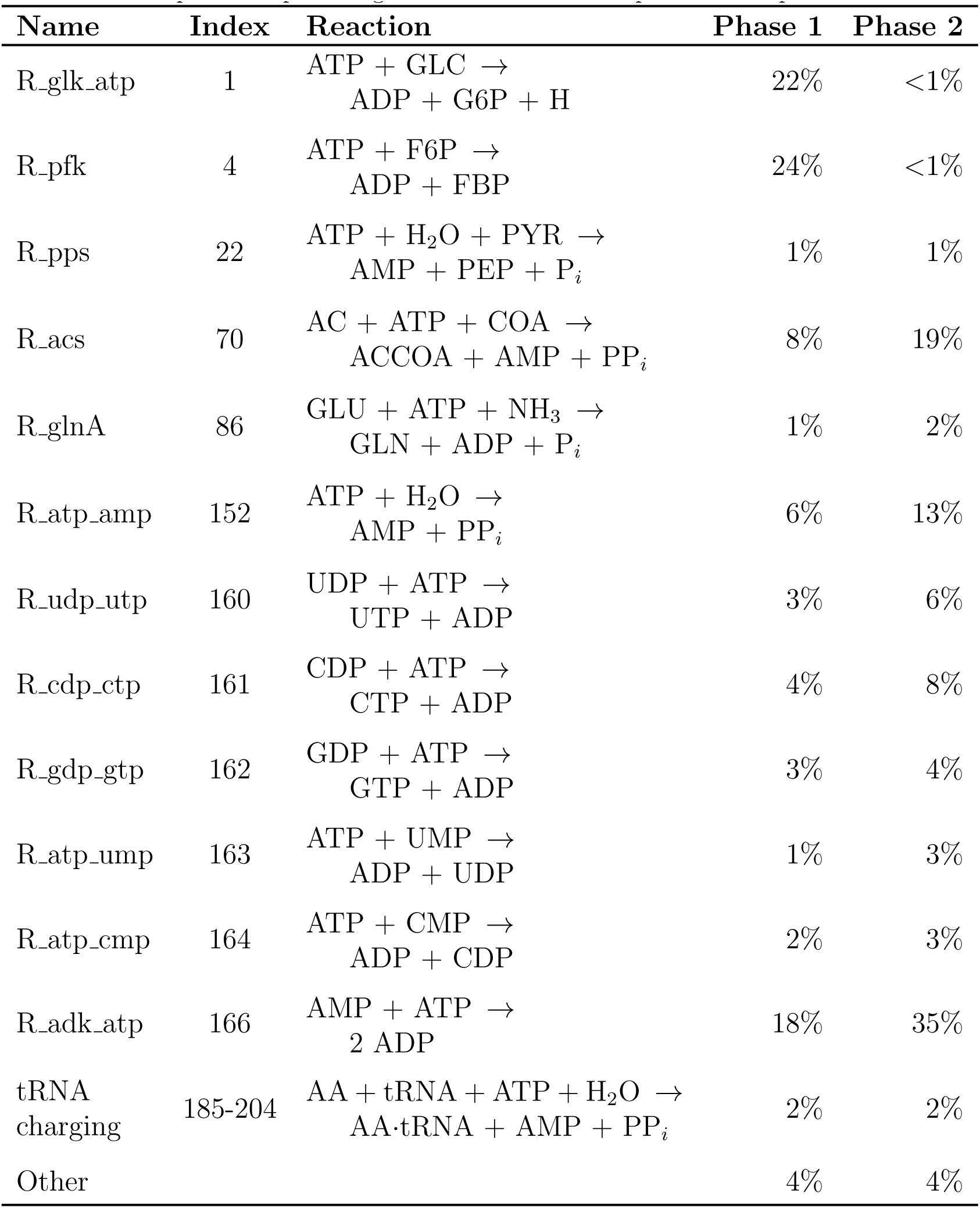
Breakdown of ATP consumption. Flux through ATP-consuming pathways in the first and second phases as percentages of total ATP consumption in that phase.

**Table 3:** Mean and standard deviation of Akaike information criterion (AIC), by measurement, for the ensemble and random ensemble.

| Measurement | $\mu_{\text{AIC}}^{\text{Ens}}$ | $\sigma_{\text{AIC}}^{\text{Ens}}$ | $\mu_{\text{AIC}}^{\text{Rand}}$ | $\sigma_{\text{AIC}}^{\text{Rand}}$ | $\mu_{\text{AIC}}^{\text{Rand}} - \mu_{\text{AIC}}^{\text{Ens}}$ |
| --- | --- | --- | --- | --- | --- |
| GLC | 65.4 | 2.1 | 103.9 | 0.6 | 38.5 |
| CAT | -23.0 | 10.5 | -5.2 | <0.1 | 17.8 |
| PYR | 64.8 | 10.3 | 84.7 | 0.7 | 19.9 |
| LAC | 70.7 | 4.5 | 88.9 | <0.1 | 18.2 |
| AC | 79.4 | 6.0 | 96 | 2.1 | 16.6 |
| SUCC | 59.6 | 3.4 | 55.5 | 4.1 | -4.1 |
| MAL | 60.8 | 4.1 | 71.6 | 6.3 | 10.8 |
| ATP | 51.1 | 3.3 | 69.1 | <0.1 | 18.0 |
| ADP | 39.8 | 3.7 | 53.2 | 4.7 | 13.4 |
| AMP | 32.9 | 1.5 | 75.1 | 5.7 | 42.2 |
| GTP | 53.4 | 1.6 | 68.2 | <0.1 | 14.8 |
| GDP | 45.7 | 2.9 | 43.6 | 9.5 | -2.1 |
| GMP | 46.5 | 4.2 | 46.1 | 12.5 | -0.4 |
| CTP | 44.9 | 2.6 | 58.5 | <0.1 | 13.7 |
| CDP | 38.8 | 1.6 | 50.7 | 8.2 | 11.8 |
| CMP | 32.1 | 4.0 | 51.9 | 9.1 | 19.8 |
| UTP | 55.6 | 5.2 | 53 | <0.1 | -2.7 |
| UDP | 28.2 | 4.6 | 51.9 | 11.5 | 23.6 |
| UMP | 35.3 | 3.3 | 72.3 | 7.3 | 36.9 |
| ALA | 66.4 | 4.4 | 100.5 | 1.1 | 34.1 |
| ASN | 53.7 | 1.5 | 67.6 | 3.8 | 13.8 |
| ASP | 65.9 | 2.5 | 79.5 | <0.1 | 13.6 |
| CYS | 60.5 | 3.1 | 74 | <0.1 | 13.5 |
| GLN | 54.3 | 5.6 | 84.7 | <0.1 | 30.4 |
| GLY | 47.2 | 12.7 | 75.5 | 11.7 | 28.3 |
| HIS | 46.3 | 6.2 | 43.2 | 3.2 | -3.2 |
| ILE | 53.3 | 3.8 | 48.4 | 4.8 | -5.0 |
| LEU | 41.5 | 6.5 | 52.5 | 4.6 | 10.9 |
| LYS | 68.4 | 2.0 | 73.9 | 0.2 | 5.5 |
| MET | 55.9 | 1.0 | 57.4 | 4 | 1.5 |
| PHE | 43.4 | 5.9 | 57.7 | 8.3 | 14.3 |
| PRO | 54.4 | 2.8 | 47.9 | 6.7 | -6.5 |
| SER | 65.9 | 4.1 | 81.4 | <0.1 | 15.6 |
| THR | 28.2 | 5.5 | 63.2 | 14.9 | 35.0 |
| TRP | 31.2 | 5.7 | 79.9 | 1.4 | 48.6 |
| TYR | 39.3 | 2.0 | 36.7 | 5.4 | -2.6 |
| VAL | 51.3 | 3.1 | 55.5 | 4.6 | 4.1 |

## Discussion

In this study, we developed an ensemble of kinetic cell-free protein synthesis (CFPS) models using dynamic metabolite measurements from an early glucose powered Cytomin *E. coli* cell-free extract. We used the hybrid cell-free modeling approach of Wayman and coworkers, which integrates traditional kinetic modeling with a logic-based description of allosteric regula-tion, to describe the time evolution of the CFPS reaction. The ensemble captured dynamic metabolite measurements over 2-orders of magnitude better than random parameter sets generated in the same region of parameter space. The ensemble captured the biphasic time course of CAT production, relying on glucose during the first hour and pyruvate and lactate following glucose exhaustion. Allosteric control was essential to the description of the organic acid trajectories; without allosteric control, pyruvate, lactate, succinate, and malate were predicted to be consumed more quickly following glucose exhaustion, to power CAT synthesis. Interestingly, CAT production was robust to the removal of allosteric control; because the amino acids and energy species that are reactants for CAT synthesis were also not affected by allosteric control. We then used the ensemble of kinetic models to analyze the performance of the CFPS system, and to estimate the pathways most important to protein production. We calculated that CAT was produced with an energy efficiency of 12%, suggesting that much of the energy resources for protein synthesis were diverted to non-productive pathways. By knocking out metabolic enzymes in groups, we showed that metabolism and protein production in particular depended upon oxidative phosphorylation and glycolysis /gluconeogenesis. Using the Sobol sampling technique we demonstrated the greater importance of translation rate than transcription. Taken together, this study provides a foundation for sequence specific genome scale, dynamic modeling of cell-free *E. coli* protein synthesis.

The ensemble of models quantitatively described the dynamic time evolution of the cell-free protein production. Thus, the model could serve as a surrogate to rationally design cell-free production processes to optimize production rate and yield. In analyzing the effect of reaction groups on CAT production and the system state, the regions of metabolism associated with substrate utilization and energy generation were the most important. Oxidative phosphorylation was vital, since it provides most of the energetic needs of CFPS. While it is unknown how active oxidative phosphorylation is compared to that of in *vivo* systems, our modeling approach suggested it was critical to CFPS performance. However, the biphasic operation of CFPS highlights the ability of the system to respond to an absence of glucose. During the first phase, there is an accumulation of central carbon metabolites with the majority of flux going toward acetate and some toward pyruvate, lactate, succinate and malate. While acetate continued to accumulate as a byproduct, the other organic acids were consumed as secondary substrates after glucose was no longer available. Glutamate also served as a substrate throughout both phases, powering amino acid synthesis. These results confirm experimental findings that CAT production can be sustained by other substrates in the absence of glucose, providing alternative strategies to optimize CFPS performance. While CAT synthesis can be powered by other substrates, the productivity is significantly lower (5 *μ*M/h, as opposed to 8 *μ*M/h). This is in accordance with literature, where pyruvate provided a relatively slow but continuous supply of ATP [28]. However, the energy efficiency is slightly higher (14% as compared with 10%). Taken together, this shows CFPS can be designed towards a specified application, either requiring a slow stable energy source or faster production.

This work represents the first dynamic model of *E. coli* cell-free protein synthesis. We apply a hybrid modeling framework to capture an experimental dataset for production of a test protein, and identify system limitations and areas of improvement for production efficiency. This work could be extended through further experimentation to gain a deeper understanding of model performance under a variety of conditions. Specifically, CAT production performed in the absence of amino acids could inform the system’s ability to manufacture them, while experimentation in the absence of glucose or oxygen could shed light on the importance of those substrates. In addition, the approach should be extended to other protein products. CAT is only a test protein used for model identification; the modeling framework, and to some extent the parameter values, should be protein agnostic. An important extension of this study would be to apply its insights to other protein applications, where possible. Having captured the experimental data, we in-vestigated if CFPS performance could be further improved. We showed that the model predicts CAT production with an energy efficiency of 10% under glucose and 14% under pyruvate. The accumulation of glycolytic intermediates and byproducts such as acetate and carbon dioxide were responsible for this sub-optimal performance. If fluxes could be balanced such that intermediates were fully utilized, CAT production would increase. Knocking out sections of network metabolism revealed that glycolysis/gluconeogenesis and oxidative phosphorylation were the most important to CAT production and the system as a whole. Productivity was also heavily dependent on the synthesis reactions of alanine, aspartate, asparagine, glutamate, and glutamine, while TCA cycle and overflow reactions affected the system state. Taken together, these findings represent the first dynamic model of *E. coli* cell-free protein synthesis, an important step toward a functional genome scale description of cell free systems.

## Materials and Methods

### Formulation and solution of the model equations

We used ordinary differential equations (ODEs) to model the time evolution of metabolite (*x_i_*), scaled enzyme activity (*ϵ_i_*), transcription *(m)* and translation 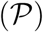 in an *E. coli* cell free metabolic network:

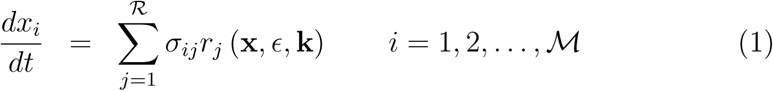

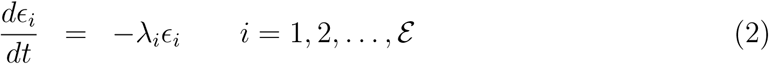

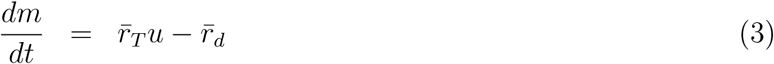

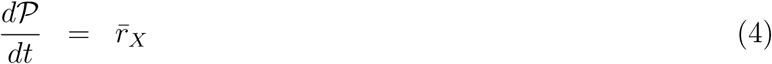

The quanity 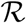 denotes the number of metabolic reactions, 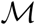 denotes the number of metabolites and 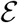 denotes the number of metabolic enzymes in the model. The quantity *r_j_*(**x**, ϵ, **k**) denotes the rate of reaction *j*. Typically, reaction *j* is a non-linear function of metabolite and enzyme abundance, as well as unknown kinetic parameters 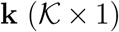. The quantity σ_*ij*_ denotes the stoichiometric coefficient for species *i* in reaction *j.* If *σ_ij_ >* 0, metabolite *i* is produced by reaction *j*. Conversely, if *σ_ij_ <* 0, metabolite i is consumed by reaction *j*, while *σ_ij_* = 0 indicates metabolite i is not connected with reaction *j*. Lastly, *λ_i_* denotes the scaled enzyme activity decay constant. The system material balances were subject to the initial conditions **x** *(t_o_)* = **x**_o_ and ϵ (*t_o_*) = **1** (initially we have 100% cell-free enzyme activity).

Metabolic reaction rates were written as the product of a kinetic term 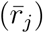 and a control term 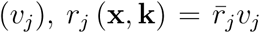. We used multiple saturation kinetics to model the reaction term 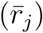:

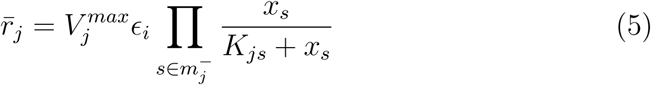

where 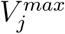 denotes the maximum rate for reaction *j*, *ϵ_i_* denotes the scaled enzyme activity which catalyzes reaction *j*, *K_js_* denotes the saturation constant for species *s,* in reaction *j* and 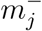 denotes the set of *reactants* for reaction *j*.

The control term 0 ≤ *V_j_* ≤ 1 depended upon the combination of factors which influenced rate process *j*. For each rate, we used a rule-based approach to select from competing control factors. If rate *j* was influenced by 1,…, *m* factors, we modeled this relationship as 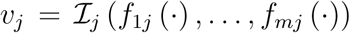 where 0 ≤ *f_ij_* (⋅) ≤ 1 denotes a transfer function quantifying the influence of factor *i* on rate *j*. The function 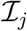 (⋅) is an integration rule which maps the output of regulatory transfer functions into a control variable. We used hill-like transfer functions and 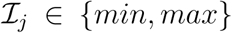 in this study [21]. We included 17 allosteric regulation terms, taken from literature, in the CFPS model. PEP was modeled as an inhibitor for phosphofructokinase [29, 30], PEP carboxykinase [29], PEP synthetase [29, 31], isocitrate dehydrogenase [29, 32], and isocitrate lyase/malate synthase [29, 32, 33], and as an activator for fructose-biphosphatase [29, 34, 35, 36]. AKG was modeled as an inhibitor for citrate synthase [29, 37, 38] and isocitrate lyase/malate synthase [29, 33]. 3PG was modeled as an inhibitor for isocitrate lyase/malate synthase [29, 33]. FDP was modeled as an activator for pyruvate kinase [29, 39] and PEP carboxylase [29, 40]. Pyruvate was modeled as an inhibitor for pyruvate dehydrogenase [29, 41, 42] and as an activator for lactate dehydrogenase [43]. Acetyl CoA was modeled as an inhibitor for malate dehydrogenase [29].

The symbol 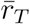 denotes the transcription rate, *u* denotes a promoter specific activation model, and 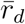 denotes the transcript degradation rate. The transcription rate was modeled as:

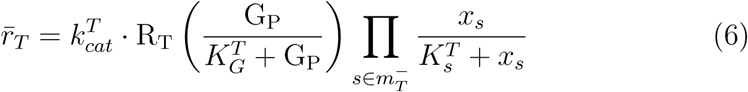

where 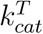 denotes the maximum transcription rate, R_T_ denotes the RNA polymerase concentration, *G_P_* denotes the gene concentration, 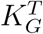 denotes the gene saturation constant, 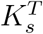 denotes the saturation constant for species s, and 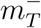 denotes the set of *reactants* for transcription: ATP, GTP, CTP, UTP, and water. In this study, we considered only the T7 promoter; we have previously estimated *u* ≃0.95 for a T7 [REF-MIKE]. While transcription was modeled as saturating with respect to gene concentration, the gene was not considered a reactant in the stoichiometry as it was not consumed. Transcript degradation was modeled as first-order in transcript:

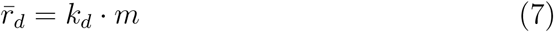

where *k^d^* denotes the transcript degradation rate constant.

The symbol 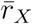 denotes the translation rate, which was modeled as:

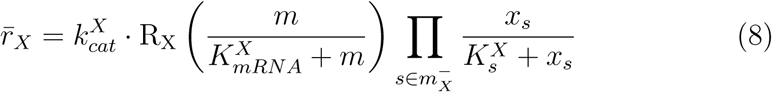

where 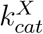 denotes the maximum translation rate, R_X_ denotes the ribosome concentration, m denotes the transcript concentration, 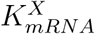 denotes the transcript saturation constant, 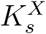 denotes the saturation constant for species s, and 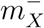 denotes the set of *reactants* for translation: GTP, water, and the 20 species representing tRNA charged with amino acids. While translation was modeled as saturating with respect to transcript concentration, the transcript was not considered a reactant in the stoichiometry as it is not consumed.

### Estimation of kinetic model parameters

We estimated an ensemble of kinetic parameter sets using a constrained Markov Chain Monte Carlo (MCMC) random walk strategy. We have used this technique previously to estimate numerically stable low-error parameter sets for signal transduction models [44, 45]. Starting from a small number of parameter sets estimated by inspection and literature, we calculated the cost function, equal to the sum-squared-error between experimental data and model predictions:

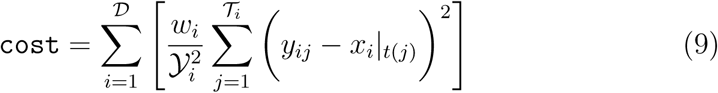

where 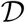 denotes the number of datasets 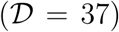, *w_i_* denotes the weight of the *i^th^* dataset, 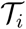 denotes the number of timepoints in the *i^th^* dataset, *t(j)* denotes the *j^th^* timepoint, *y_ij_* denotes the measurement value of the *i^th^* dataset at the *j^th^* timepoint, and x_i_|_t(j)_ denotes the simulated value of the metabolite corresponding to the *i^th^* dataset, interpolated to the *j^th^* timepoint. Lastly, the cost function was scaled by the maximum experimental value in the *i^th^* dataset, 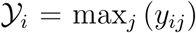. We then perturbed each model parameter between an upper and lower bound that varied by parameter type:

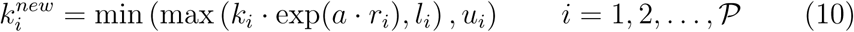

where 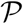 denotes the number of parameters 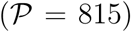, which includes 204 maximum reaction rates (*V^max^*), 204 enzyme activity decay constants, 548 saturation constants (*K_js_*), and 34 control parameters, 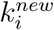 denotes the new value of the *i^th^* parameter, *k_i_* denotes the current value of the *i^th^* parameter, *a* denotes a distribution variance, *r_i_* denotes a random sample from the normal distribution, *l_i_* denotes the lower bound for that parameter type, and *u_i_* denotes the upper bound for that parameter type. Model parameters were constrained by literature collected using the BioNumbers database [22]. Transcription, translation, and mRNA degradation were bounded within a factor of two of their reference values. A characteristic cell-free enzyme concentration of 170 nM was calculated by diluting the one-tenth maximal con-centration of *lacZ* (5 *μ*M, BNID 100735) by a cell-free dilution factor of 30. This enzyme level was then used to calculate rate maxima from turnover numbers for various enzymes from BioNumbers (Table 4). Rate maxima were bounded within one order of magnitude of the reference value where available; all other rate maxima were bounded within two orders of magnitude of the geometric mean of the available values. Enzyme activity decay constants were bounded between 0 and 1 h^−1^, corresponding to half lives of 42 minutes and infinity. Saturation constants were bounded between 0.0001 and 10 mM. Control gain parameters were bounded between 0.05 and 10 (dimensionless), while order parameters were bounded between 0.02 and 10 (dimensionless).

For each newly generated parameter set, we re-solved the balance equations and calculated the cost function. All sets with a lower cost were accepted into the ensemble. Sets with a higher cost were also accepted into the ensemble, if they satisfied the acceptance constraint:

**Table 4:** Reference values for reaction rate maxima (V_*max*_) from BioNumbers. *V_max_* values calculated from turnover numbers (k_*cat*_) from BioNumbers, and a characteristic enzyme concentration of 170 nM. Characteristic rate maximum for all other reactions calculated as geometric mean of calculated rate maxima.

| Enzyme | Reaction | $k_{cat}$ ($\text{min}^{-1}$) | $V_{max}$ (mM/h) | BNID# |
| --- | --- | --- | --- | --- |
| Serine dehydrase | R_ser_deg | 10400 | 104 | 101119 |
| Isocitrate dehydrogenase | R_icd | 11900 | 119 | 101152 |
| Lactate dehydrogenase | R_ldh | 5800 | 58 | 101036 |
| Aspartate transaminase | R_aspC | 25800 | 258 | 101108 |
|  | R_tyr |  |  |  |
|  | R_phe |  |  |  |
| Enolase | R_eno | 13200 | 132 | 101028 |
| Pyruvate kinase | R_pyk | 25000 | 250 | 101029 |
|  |  |  |  | 101030 |
| Malic enzyme | R_maeA | 35400 | 354 | 101167 |
|  | R_maeB |  |  |  |
| Phosphofructokinase | R_pfk | 554400 | 5544 | 104955 |
| Malate dehydrogenase | R_mdh | 33000 | 330 | 101163 |
| Citrate Synthase | R_gltA | 42000 | 420 | 101149 |
| 6PG dehydrogenase | R_zwf | 3200 | 32 | 101048 |
|  | R_pgl |  |  |  |
|  | R_gnd |  |  |  |
| Succinate dehydrogenase | R_sdh | 121 | 1.21 | 101162 |
| Succinyl-coA synthetase | R_sucCD | 4700 | 47 | 101158 |
| 3PGA dehydrogenase | R_gpm | 1100 | 11 | 101135 |
| PEP carboxylase | R_ppc | 35400 | 354 | 101139 |
| 3PGA kinase | R_pgk | 4300 | 43 | 101016 |
| Characteristic $V_{max}$ | | | 110 | |

**Table 5:** Reference values for transcription, translation, and mRNA degradation from literature. Transcription rate calculated from elongation rate, mRNA length, and promoter activity level. Translation rate calculated from elongation rate, protein length, and polysome amplification constant. mRNA degradation rate calculated from mRNA degradation time.

| Description | Parameter | Value | Units | Reference |
| --- | --- | --- | --- | --- |
| T7 RNA polymerase concentration | $R_T$ | 1.0 | $\mu\text{M}$ | |
| Ribosome concentration | $R_X$ | 2 | $\mu\text{M}$ | [10] |
| Transcription saturation coefficient | $K_T$ | 100 | nM | estimated |
| Translation saturation coefficient | $K_X$ | 45 | $\mu\text{M}$ | estimated |
| Transcription elongation rate | $\dot{v}_T$ | 25 | nt/s | [10] |
| CAT mRNA length | $l_G$ | 660 | nt | [48] |
| Promoter activity level | $u$ | 0.9 | | estimated |
| Transcription rate | $k_{cat}^T = \left(\frac{\dot{v}_T}{l_G}\right) u$ | 123 | $\text{h}^{-1}$ | calculated |
| Translation elongation rate | $\dot{v}_X$ | 1.5 | aa/s | [10] |
| CAT protein length | $l_P$ | 219 | aa | [48] |
| Polysome amplification constant | $K_P$ | 10 | | estimated |
| Translation rate | $k_{cat}^X = \left(\frac{\dot{v}_X}{l_P}\right) K_P$ | 247 | $\text{h}^{-1}$ | calculated |
| mRNA degradation time | $t_{1/2}$ | 8 | min | BNID 106253 |
| mRNA degradation rate | $k_{deg} = \frac{\ln(2)}{t_{1/2}}$ | 5.2 | $\text{h}^{-1}$ | calculated |
| ATP transcription coefficient | $\text{ATP}_T$ | 176 | | calculated |
| CTP transcription coefficient | $\text{CTP}_T$ | 144 | | calculated |
| GTP transcription coefficient | $\text{GTP}_T$ | 151 | | calculated |
| UTP transcription coefficient | $\text{UTP}_T$ | 189 | | calculated |
| ATP tRNA charging coefficient | $\text{ATP}_X$ | 219 | | calculated |
| GTP translation coefficient | $\text{GTP}_X$ | 438 | | calculated |

**Table 6:**
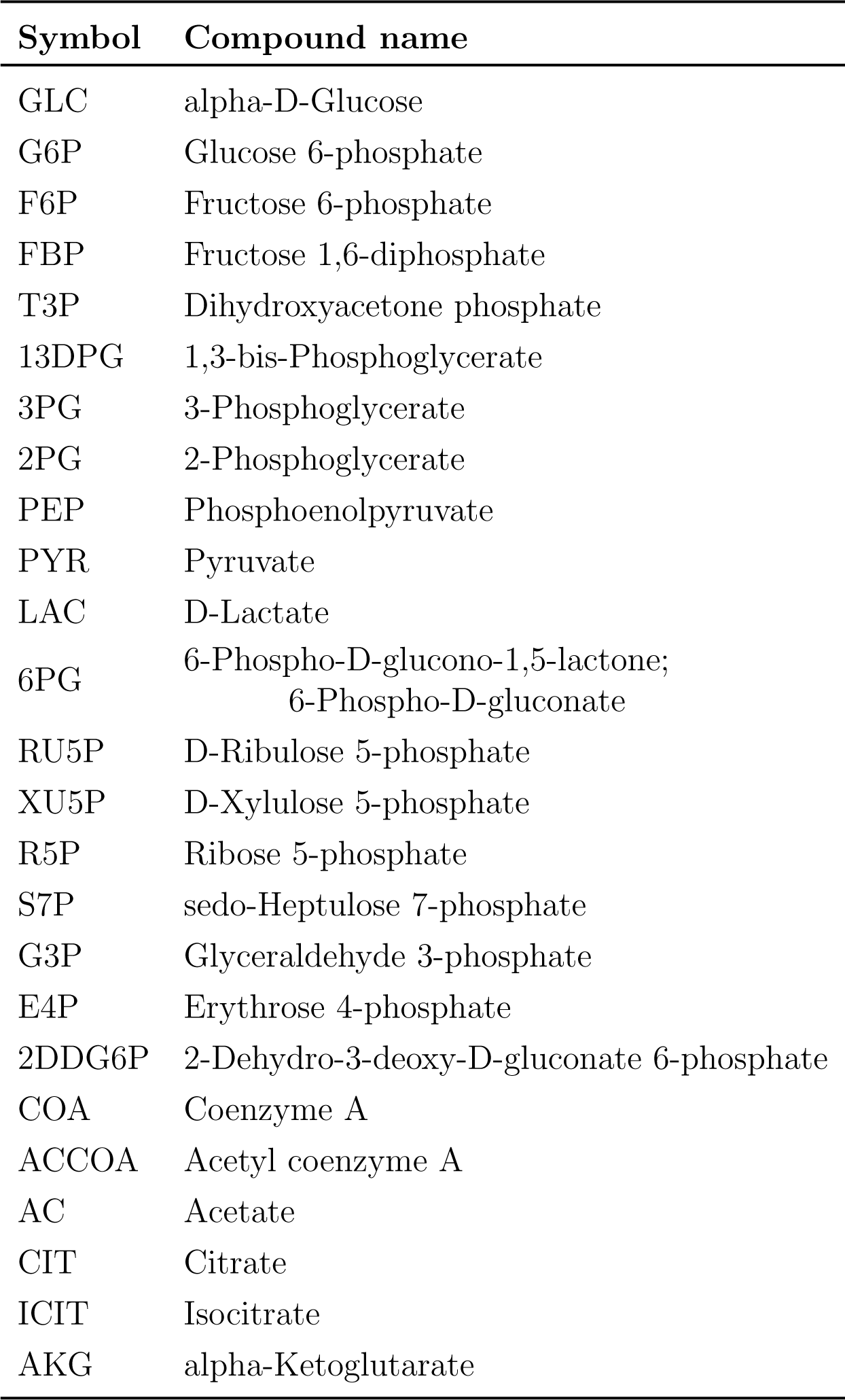
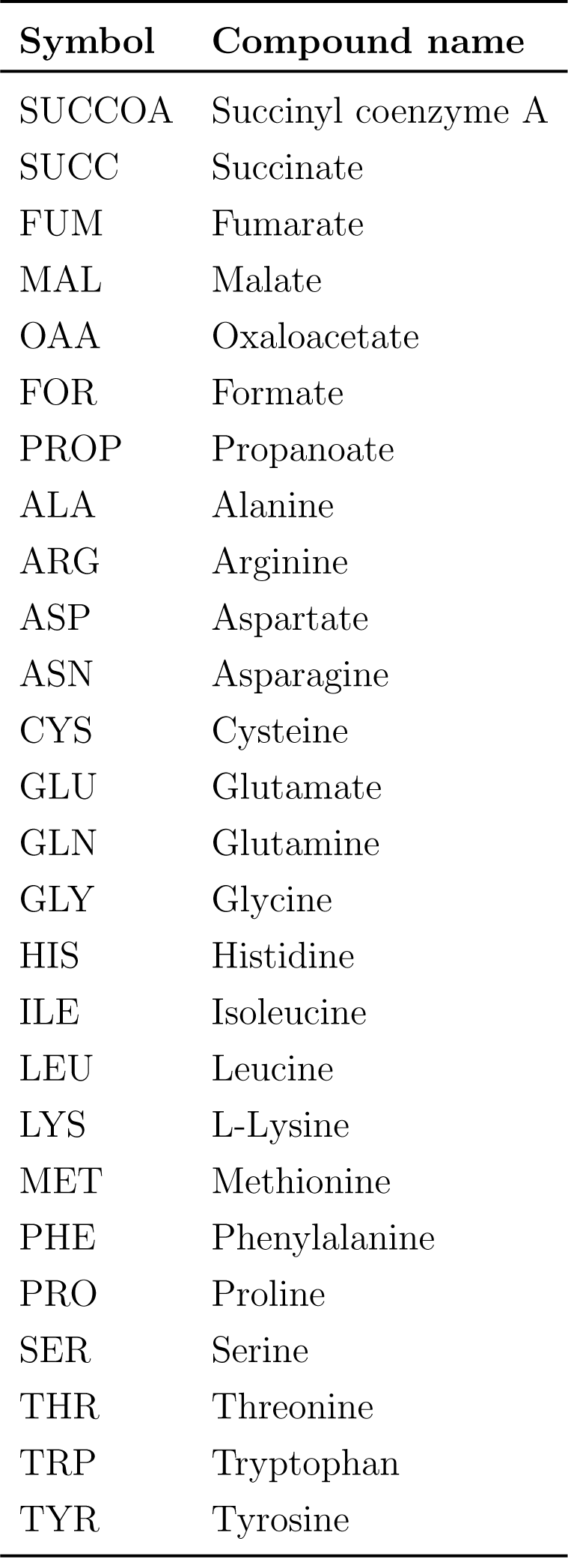
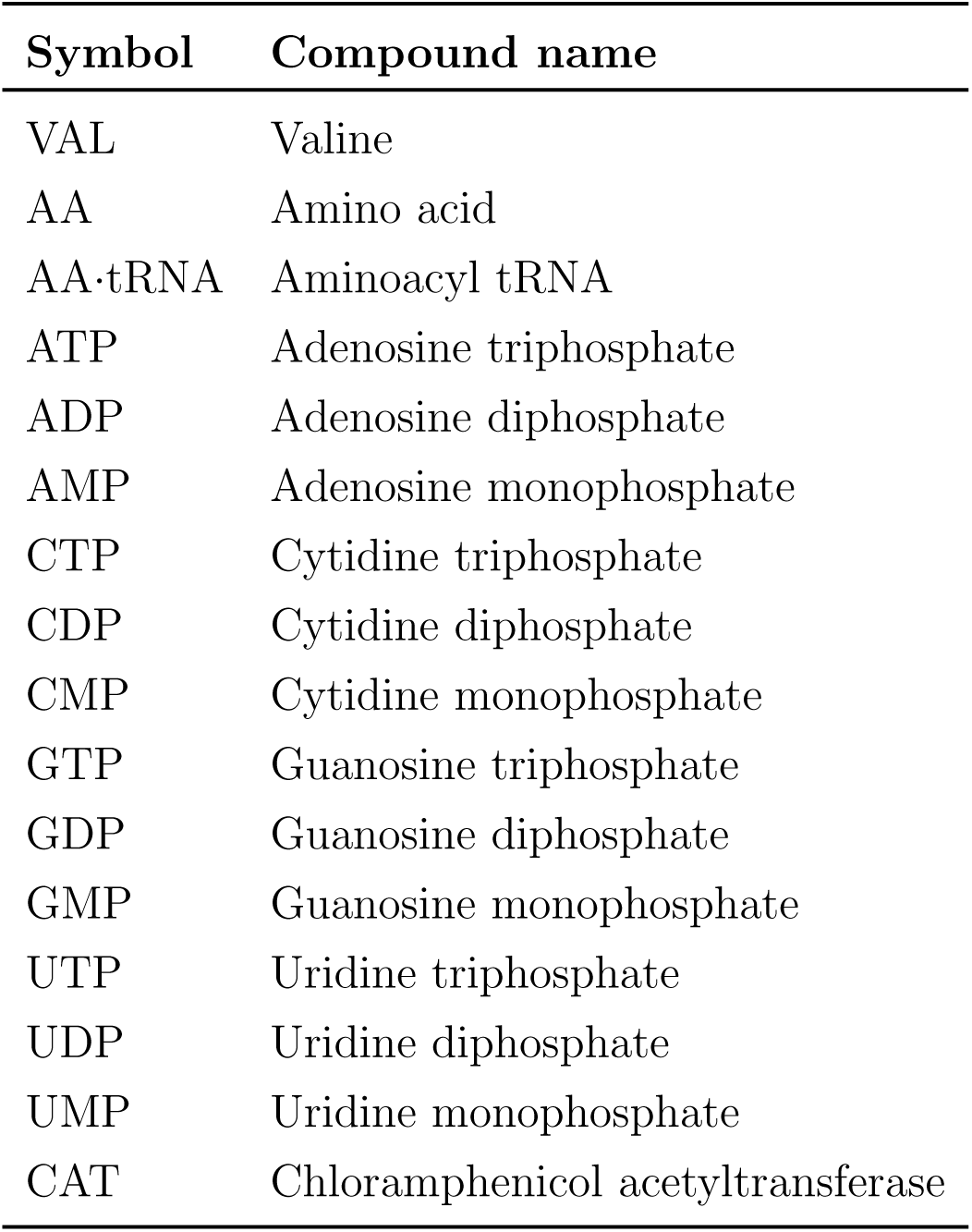
Nomenclature

| Symbol | Compound name |
| --- | --- |
| GLC | alpha-D-Glucose |
| G6P | Glucose 6-phosphate |
| F6P | Fructose 6-phosphate |
| FBP | Fructose 1,6-diphosphate |
| T3P | Dihydroxyacetone phosphate |
| 13DPG | 1,3-bis-Phosphoglycerate |
| 3PG | 3-Phosphoglycerate |
| 2PG | 2-Phosphoglycerate |
| PEP | Phosphoenolpyruvate |
| PYR | Pyruvate |
| LAC | D-Lactate |
| 6PG | 6-Phospho-D-glucono-1,5-lactone;<br>6-Phospho-D-gluconate |
| RU5P | D-Ribulose 5-phosphate |
| XU5P | D-Xylulose 5-phosphate |
| R5P | Ribose 5-phosphate |
| S7P | sedo-Heptulose 7-phosphate |
| G3P | Glyceraldehyde 3-phosphate |
| E4P | Erythrose 4-phosphate |
| 2DDG6P | 2-Dehydro-3-deoxy-D-gluconate 6-phosphate |
| COA | Coenzyme A |
| ACCOA | Acetyl coenzyme A |
| AC | Acetate |
| CIT | Citrate |
| ICIT | Isocitrate |
| AKG | alpha-Ketoglutarate |
| SUCCOA | Succinyl coenzyme A |
| SUCC | Succinate |
| FUM | Fumarate |
| MAL | Malate |
| OAA | Oxaloacetate |
| FOR | Formate |
| PROP | Propanoate |
| ALA | Alanine |
| ARG | Arginine |
| ASP | Aspartate |
| ASN | Asparagine |
| CYS | Cysteine |
| GLU | Glutamate |
| GLN | Glutamine |
| GLY | Glycine |
| HIS | Histidine |
| ILE | Isoleucine |
| LEU | Leucine |
| LYS | L-Lysine |
| MET | Methionine |
| PHE | Phenylalanine |
| PRO | Proline |
| SER | Serine |
| THR | Threonine |
| TRP | Tryptophan |
| TYR | Tyrosine |
| VAL | Valine |
| AA | Amino acid |
| AA·tRNA | Aminoacyl tRNA |
| ATP | Adenosine triphosphate |
| ADP | Adenosine diphosphate |
| AMP | Adenosine monophosphate |
| CTP | Cytidine triphosphate |
| CDP | Cytidine diphosphate |
| CMP | Cytidine monophosphate |
| GTP | Guanosine triphosphate |
| GDP | Guanosine diphosphate |
| GMP | Guanosine monophosphate |
| UTP | Uridine triphosphate |
| UDP | Uridine diphosphate |
| UMP | Uridine monophosphate |
| CAT | Chloramphenicol acetyltransferase |

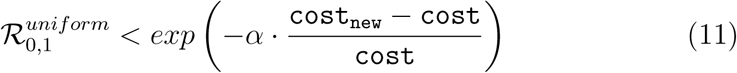

where 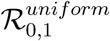 denotes a random number taken from a uniform distribution between 0 and 1, cost denotes the cost of the current parameter set, cost_new_ denotes the cost of the new parameter set, and *α* denotes a tunable parameter to control the tolerance to high-error sets. A total of 3,875 sets were accepted into the initial ensemble, from which we selected N = 100 with minimal error for the final ensemble.

Lastly, a random ensemble of 100 parameter sets was generated within the same parameter bounds as the trained ensemble. The randomized parameter sets were generated using a Monte Carlo approach: each parameter was taken from a uniform distribution constructed between its upper and lower bounds. The model equations were then solved and the cost function, and the Akaike information criterion (AIC) were calculated for each of the 37 separate experimental datasets.

### Reaction group knockouts

The metabolic network was divided into 19 reaction groups: glycolysis/gluconeogenesis, pentose phosphate, Entner-Doudoroff, TCA cycle, oxidative phosphorylation, cofactor reactions, anaplerotic/glyoxylate reactions, overflow metabolism, folate synthesis, purine/pyrimidine reactions, alanine/ aspartate/asparagine synthesis, glutamate/glutamine synthesis, arginine/proline synthesis, glycine/serine synthesis, cysteine/methionine synthesis, threonine/ lysine synthesis, histidine synthesis, tyrosine/tryptophan/phenylalanine synthesis, and valine/leucine/isoleucine synthesis. Each reaction group and pair of reaction groups were removed and the model was re-solved; the CAT productivity was then calculated and subtracted from that of the base case (no knockouts):

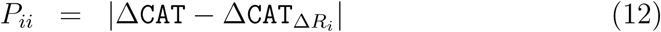

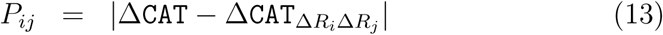

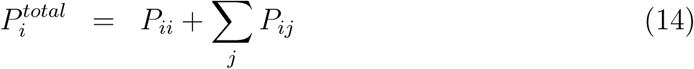

where *P_ii_* denotes the first-order productivity knockout effect for reaction group *i, P_ij_* denotes the pairwise productivity knockout effect for reaction groups *i* and *j*, 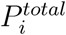 denotes the total-order productivity knockout effect for reaction group *i,* `ΔCAT` denotes the base case CAT productivity, `ΔCAT`_ΔRi_ denotes the CAT productivity when reaction group i is knocked out, `ΔCAT`_ΔRiΔRj_ denotes the CAT productivity when reaction groups i and j are knocked out, and |x| denotes the absolute value of x. The system state, defined as the model predictions for all species for which experimental data exists, was also recorded for each knockout and compared to the base case:

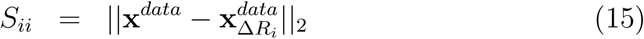

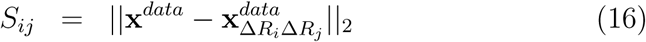

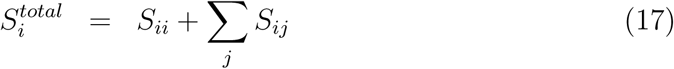

where *S_ii_* denotes the first-order system state knockout effect for reaction group *i, Sij* denotes the pairwise system state knockout effect for reaction groups i and j, 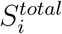 denotes the total-order system state knockout effect for reaction group i, *x^data^* denotes the base-case system state, 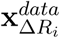 denotes the system state when reaction group i is knocked out, 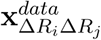 denotes the system state when reaction groups i and j are knocked out, and ||x*||_2_* denotes the *l*^*2*^ norm of x. In order to not dominate the colorbar, the total-order knockout effects were normalized to the same ranges as the main arrays (first-order and pairwise effects).

### Sensitivity of CAT productivity to transcription and translation

The catalytic rates of transcription and translation were sampled within one order of magnitude on each side from the best-fit values. The parameter bounds were set as the base*-10* logarithms of the upper and lower bound for each rate; then, *10* was taken to the power of each parameter sample to obtain the catalytic rates:

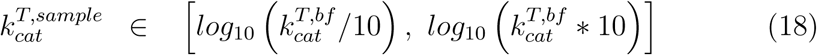

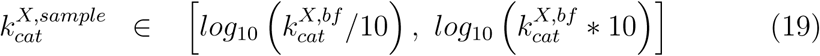

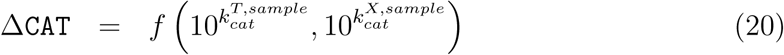

where 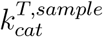 denotes the sample of the transcription catalytic rate, 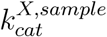 denotes the sample of the translation catalytic rate, 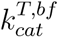 denotes the best-fit value of the transcription catalytic rate, and 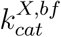 denotes the best-fit value of the translation catalytic rate. The sampling was performed using the Sensitivity Analysis Library in Python (Numpy) with 3000 samples [46].

### Calculation of energy efficiency

Energy efficiency was calculated as the ratio of transcription and translation (weighted by the appropriate energy species coefficients) to ATP generation:

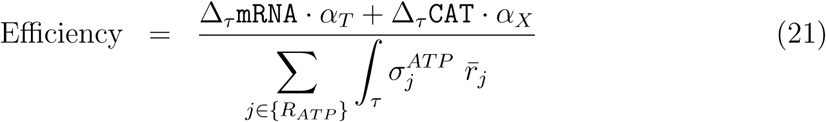

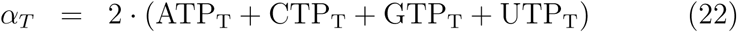

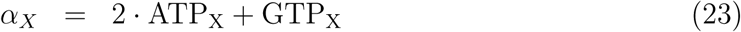

where Δ_T_`mRNA` denotes the net accumulation of mRNA in phase τ (first, second, or overall), Δ_T_`CAT` denotes the net accumulation of protein in phase τ, *α_T_* denotes the energy cost of transcription, *α_x_* denotes the energy cost of translation, *R_ATP_* denotes the set of ATP-producing reactions, and 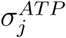 denotes the ATP coefficient for reaction *j*. ATP_T_, CTP_T_, GTP_T_, UTP_T_ denote the stoichiometric coefficients of each energy species for transcription, and ATP_X_, GTP_X_ denote the stoichiometric coefficients of ATP and GTP for translation. During transcription and tRNA charging, triphosphate molecules are consumed with monophosphates as byproducts; this is the reason for the factors of 2 on ATP_T_, CTP_T_, GTP_T_, UTP_T_, and ATP_X_.

### Availability of model code

The cell free model equations, and the parameter estimation procedure, were implemented in the Julia programming language. The model equations were solved using the **`CVODE`** solver of the **`SUNDIALS`** suite [47], with an absolute tolerance and relative tolerance of 1e^−9^; any sets exhibiting **`CVODE`**errors were discarded. Thus, the numerical stability of all parame-ters in the ensemble was ensured. The model code and parameter ensemble is freely available under an MIT software license and can be downloaded from <u>http://www.varnerlab.org</u>.

## Competing interests

The authors declare that they have no competing interests.

## Author’s contributions

J.V directed the modeling study. K.C and J.S conducted the cell-free protein synthesis experiments. J.V, J.W, and N.H developed the cell-free protein synthesis mathematical model, and parameter ensemble. The manuscript was prepared and edited for publication by J.S, N.H, M.V, J.W and J.V.

## Acknowledgements

We gratefully acknowledge the suggestions from the anonymous reviewers to improve this manuscript.

## Funding

This study was supported by a National Science Foundation Graduate Research Fellowship (DGE-1333468) to N.H. Research reported in this publication was also supported by the Systems Biology Coagulopathy of Trauma Program with support from the US Army Medical Research and Materiel Command under award number W911NF-10-1-0376.

---

**Figure S1:**
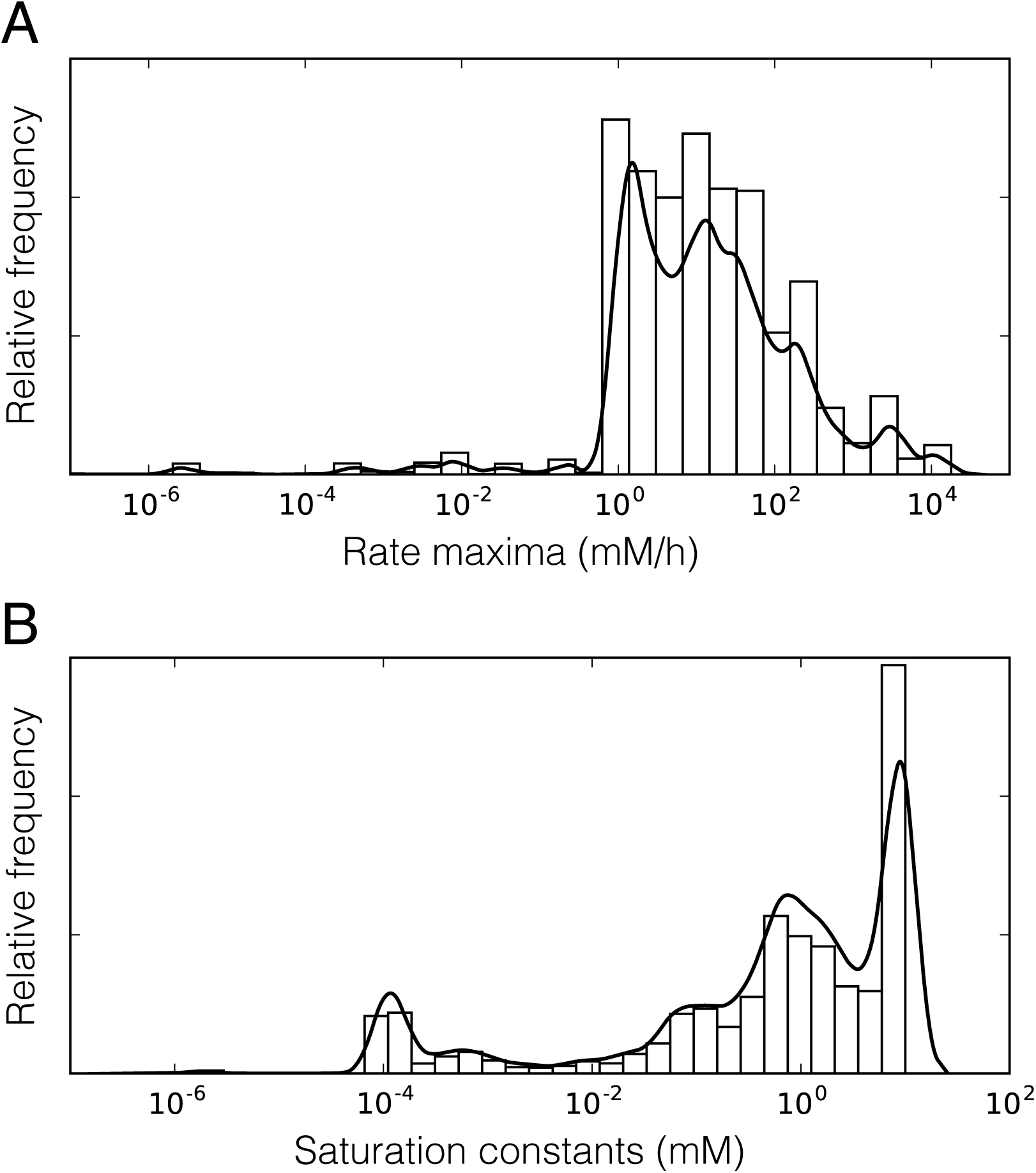
Histograms of model parameters, across the ensemble of 100 sets. A. Histogram of rate maxima. B. Histogram of saturation constants.

